# Cognitive Load modulates the exploration-exploitation trade-off Through Reconfiguration of an IPS-Centered Decision Network

**DOI:** 10.64898/2026.11.21.768355

**Authors:** Haowei Dai, Linhui Wang, Xueping Yin, Xiangyi Liang, Yutong Ying, Yuanyuan Zeng, Lijing Niu, Jonathan P. Roiser, Xiangliang Tan, Ruibin Zhang

**Affiliations:** Cognitive Control and Brain Healthy Laboratory, Department of Psychology, School of Public Health, Southern Medical University, Guangzhou, China; Department of Medical Imaging Center, Nanfang Hospital, Southern Medical University, Guangzhou, China; Institute of Cognitive Neuroscience, University College London, London, UK

**Keywords:** Cognitive load, Directed exploration, Exploration–exploitation trade-off, Dynamic decision-making, Functional connectivity

## Abstract

In dynamic environments where rewards continuously change, adaptive decision-making requires ongoing updates of option-value and uncertainty estimates to balance exploration and exploitation. This process depends on cognitive resources and involves coordinated interactions among multiple brain regions; however, how cognitive load affects directed exploration in such dynamic environments, and whether these effects are accompanied by changes in interactions among exploration-related brain regions, remains unclear. We embedded a working-memory load manipulation within a dynamic four-armed bandit task. Experiment 1 compared computational models and quantified load-related changes in exploration; Experiment 2 examined these behavioral effects in an independent sample and investigated their neural correlates using fMRI. Across both experiments, high load reduced the uncertainty bonus parameter (φ), indicating weaker directed exploration. Experiment 2 additionally showed lower inverse temperature (β) under high load, indicating more stochastic choices. Directed exploration relative to exploitation more activated the frontopolar cortex (FP), intraparietal sulcus (IPS), anterior insula (aIns), and dorsal anterior cingulate cortex (dACC). Psychophysiological interaction (PPI) analyses indicated stronger functional coupling between the right aIns and IPS under low load. Guided by the candidate connections identified by PPI, dynamic causal modeling (DCM) focused on a three-region subnetwork. Bayesian model selection favored the model with directed-exploration-related input to the IPS, while condition-specific comparisons suggested stronger aIns-to-IPS effective connectivity under low load. Together, these findings link cognitive resource demands to reduced uncertainty-guided directed exploration in a dynamic environment and suggest that this behavioral change is accompanied by selective reconfiguration of interactions within an IPS-centered decision subnetwork.

## 1. Introduction

In dynamic environments with incomplete information, adaptive decision-making requires individuals to balance the exploitation of currently valuable options against the exploration of uncertain alternatives, thereby trading off immediate reward against future information gain. This exploration–exploitation (E–E) trade-off is a fundamental component of adaptive decision-making (Cohen et al., 2007; Daw et al., 2006; Lloyd et al., 2025). Compared with relatively stable environments, previously learned values may become outdated as rewards change, requiring individuals to continuously update estimates of expected value and uncertainty based on feedback (Bond et al., 2021). Human exploration, however, is not a unitary process and can be decomposed into at least two computationally distinct strategies. Directed exploration (DE) assigns additional information value to uncertain or information-rich options, systematically biasing choice toward options that can reduce uncertainty, whereas random exploration (RE) mainly reflects a weaker influence of value on choice, expressed as increased choice stochasticity or variability (Gershman, 2018; Harms et al., 2024; Wilson et al., 2014). In dynamic environments, DE requires continuous updating of uncertainty estimates and integration of potential information gains into decision values, whereas RE does not rely on information value in the same way. These decisions involve multiple cognitive functions, including working memory, attentional allocation, and cognitive control, and may therefore depend on both limited cognitive resources and coordinated interactions among brain regions (Wyatt et al., 2024). This raises an important question: How does higher cognitive load, which increases demands on limited cognitive resources, affect directed and random exploration and the network interactions supporting these strategies in dynamic environments?

Behavioral and computational studies have shown that cognitive demands influence exploration strategies, but these effects may vary across forms of exploration and task contexts. Cogliati Dezza et al. found that increasing concurrent cognitive load reduced directed exploration while increasing random exploration, and their computational modeling further suggested that this shift was associated with impaired integration of reward information and information value (Cogliati Dezza et al., 2019). Similarly, increasing memory demands in a multi-armed bandit task reduced strategic exploration of unknown options and increased reliance on existing information (Brown et al., 2022), whereas higher working memory capacity was associated with greater directed exploration (Duan et al., 2026). Moreover, recent research reported that working memory load increased information sampling in a four-armed bandit task with relatively stable rewards, further suggesting that load effects may depend on task structure and specific components of exploration (Wan & Sloutsky, 2026). Although these studies have established a relationship between cognitive demands and exploratory behavior, the effects of cognitive load may depend on the dynamics of the reward environment. For example, the Horizon Task was specifically designed to distinguish directed from random exploration by manipulating information availability and future opportunities for exploration, while reward distributions remain stable within each game (Wilson et al., 2014). In contrast, dynamic four-armed bandit tasks allow option values to change continuously, requiring individuals to repeatedly update their estimates of expected value and uncertainty (Daw et al., 2006; Wiehler et al., 2021). This distinction raises the question of how cognitive load affects uncertainty-guided exploration when reward values continuously change. Furthermore, whether these effects are accompanied by changes in coordination among exploration-related brain regions remains unclear.

Cognitive resources support decision-making through several related processes, including working memory, attentional allocation, and cognitive control, which depend on coordination across distributed brain regions (Cole et al., 2013; Duncan, 2010). Therefore, when cognitive load increases demand on limited cognitive resources, its neural effects may involve not only changes in regional activity but also adjustments in interactions between regions. Previous studies have shown that working memory load can modulate functional and effective connectivity among frontal, parietal, and other task-related regions (Aben et al., 2020; Turker et al., 2026; Zhao et al., 2024). In exploration–exploitation decisions, neuroimaging studies have identified several exploration-related regions, including the frontopolar cortex (FP), intraparietal sulcus (IPS), anterior insula (aIns), and dorsal anterior cingulate cortex (dACC) (Daw et al., 2006; Wiehler et al., 2021). Recent neuroimaging evidence has further implicated attentional, cognitive-control, and salience-related networks in exploratory decisions (Wyatt et al., 2024), A meta-analysis of 23 studies also highlighted the involvement of the aIns and dACC, while revealing substantial overlap in regional activity between exploration and exploitation (Sazhin et al., 2025). However, conventional univariate activation analyses mainly characterize regional responses to task conditions and do not directly reveal how cognitive load changes coordination between these regions. Graph-theoretical approaches characterize network topology, such as network integration and node centrality, but these measures alone do not directly specify task-dependent coupling or directional influences between particular regions (Bullmore & Sporns, 2009). In contrast, psychophysiological interaction (PPI) analysis examines condition-dependent changes in functional coupling (O’Reilly et al, 2012), while dynamic causal modeling (DCM) compares neural network models to evaluate driving-input locations and estimate directional effective connectivity (Friston et al., 2003). Combining regional activation, PPI, and DCM analyses may therefore help determine whether cognitive load is accompanied by selective reconfiguration of interactions within exploration-related networks, providing a network-level account of how cognitive resources influence directed exploration.

To further understand how cognitive load may alter network interactions, it is important to consider the computational roles of different exploration-related regions. Among these regions, the IPS is of particular interest because it may contribute to representing uncertainty and using this information to guide DE. Previous studies have implicated the posterior parietal cortex in decision-making under uncertainty and short-term information maintenance (Huettel et al., 2005; Todd & Marois, 2004). In dynamic environments where rewards continuously change, these functions are relevant to updating estimates of option values and uncertainty. Recent evidence further suggests that the IPS encodes exploration signals related to currently available options (Chan et al., 2026). More direct evidence comes from Wiehler et al., who used DCM to compare different locations of DE-related driving inputs and found the strongest support for the model with input to the IPS (Wiehler et al., 2021). This finding provides a basis for further examining the role of the IPS in exploration-related network interactions. Meanwhile, the aIns is involved in uncertainty and salience processing, and its functional connectivity can vary with cognitive task demands (Cai & Menon, 2025; Menon & Uddin, 2010). The dACC contributes to evaluating the potential benefits of behavioral choices and the costs and benefits of cognitive control (Crawford et al., 2024; Kolling et al., 2012; Shenhav et al., 2013). In addition, cognitive effort demands have been shown to modulate functional connectivity between the dACC and task-relevant regions (Aben et al., 2020). These findings suggest that cognitive load may influence not only regional activity but also coordination among exploration-related regions during uncertainty processing, value evaluation, and cognitive control. However, whether cognitive load alters connections involving the IPS within the exploration-related network, and whether these changes are associated with reduced DE, remains unclear.

Building on this framework, we proposed that higher cognitive load may constrain the use of uncertainty information in dynamic environments, thereby reducing directed exploration, and may be accompanied by changes in coordination among exploration-related brain regions, particularly interactions involving the IPS. To examine this possibility, we used a dynamic four-armed bandit task with a concurrent working memory task to manipulate cognitive load across two experiments. Experiment 1 used computational model comparison and parameter estimation to examine the effects of cognitive load on DE and RE. Experiment 2 further examined these behavioral effects in an independent sample and combined the task with fMRI to investigate their neural basis.

Specifically, we first examined regional activity in exploration-related regions, including the FP, IPS, aIns, and dACC, and then used PPI to assess load-related changes in task-dependent functional coupling among these regions. Based on the candidate connections identified by PPI, we defined a subnetwork for DCM and further compared different locations of DE-related driving inputs and effective-connectivity parameters during DE under high and low load (Figure 1). By combining computational modeling, regional activation, and network connectivity analyses, we aimed to investigate whether reduced DE under cognitive load is accompanied by selective reconfiguration of connectivity patterns within exploration-related networks, particularly those involving the IPS, providing a network-level account of how cognitive resources influence directed exploration in dynamic environments.

**Figure 1.**
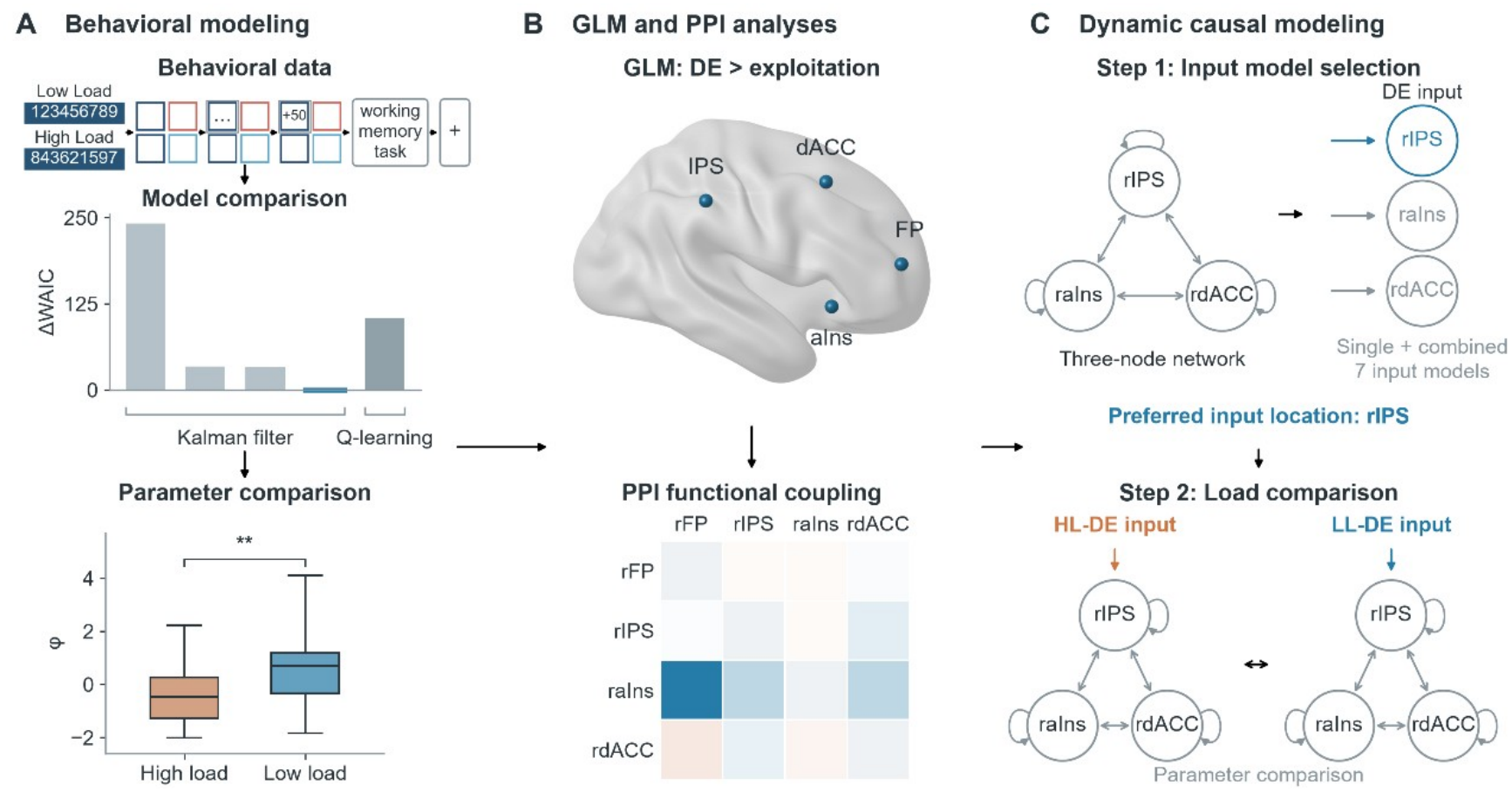
Overview of the experimental task and analysis pipeline. (A) Behavioral modeling. Participants completed a dynamic four-armed bandit task with a concurrent working memory task under high- and low-cognitive-load conditions. Choice and reward data were fitted using Q-learning and Kalman filter models, which were compared using the Watanabe–Akaike information criterion (WAIC). Model parameters were then compared between load conditions. (B) GLM and PPI analyses. General linear model (GLM) analyses examined brain activity associated with directed exploration relative to exploitation. Four exploration-related regions were examined: the frontopolar cortex (FP), intraparietal sulcus (IPS), anterior insula (aIns), and dorsal anterior cingulate cortex (dACC). Psychophysiological interaction (PPI) analyses further examined load-related changes in functional coupling among these regions. (C) Dynamic causal modeling. Dynamic causal modeling (DCM) was performed using a three-node network comprising the rIPS, raIns, and rdACC. In Step 1, seven candidate models with different directed-exploration driving-input locations were compared, identifying rIPS as the preferred input location. In Step 2, condition-specific DCMs for high-load and low-load directed exploration were compared to assess load-related differences in effective connectivity. Abbreviations: DE, directed exploration; HL, high load; LL, low load; ROI, region of interest.

## 2. Methods

### 2.1 Samples

The study included two experiments. In Experiment 1, 27 healthy participants were recruited for the behavioral experiment. All participants were screened using the Generalized Anxiety Disorder-7 (GAD-7) and Patient Health Questionnaire-9 (PHQ-9), and only individuals with GAD-7 scores < 5 and PHQ-9 scores < 5 were included.

In Experiment 2, 40 participants completed the combined cognitive-load and decision-making task and were included in the behavioral analyses. Ten participants did not show any directed exploration trials and therefore could not be included in the subsequent imaging analyses based on directed-exploration events. The final fMRI sample therefore included 30 participants. We further confirmed that exclusion of these participants did not change the main behavioral results (Supplementary Results 1).

All participants provided written informed consent before participation. The study was approved by the Biomedical Ethics Committee of Southern Medical University (approval no. Nan Yi Lun Li Shen [2026] No. 35) and was conducted in accordance with the approved ethical procedures.

### 2.2 Task and procedure

We combined a dynamic four-armed bandit task with a concurrent working memory task to manipulate cognitive load, drawing on established procedures (Cogliati Dezza et al., 2019; Daw et al., 2006). On each trial, participants selected one of four colored options and received a reward ranging from 0 to 100 points. Reward values for the four options evolved independently according to Gaussian random walks, causing option values to change throughout the experiment.

Participants were instructed to earn as many points as possible. The task consisted of eight blocks of 35 trials (280 trials in total), with short breaks between blocks. Final compensation included a fixed payment and a performance-based bonus.

At the beginning of each block, participants were presented with a nine-digit sequence and were required to remember it throughout the block. Each trial consisted of a choice, reward feedback, and a working memory probe. After reward feedback, one of the first eight digits in the sequence was presented as a probe. Participants had 1500 ms to press the key corresponding to the digit that followed the probe in the memorized sequence. For example, if the sequence was “123456789” and the probe was “3,” the correct response was “4.”

Cognitive load was manipulated at the block level using high- and low-load conditions. Under high load, the nine digits were presented in a randomized order (e.g., “864235179”) for 2000 ms, with a new random sequence used in each high-load block. Under low load, the fixed sequence “123456789” was presented for 500 ms. The order of high- and low-load blocks was randomized and counterbalanced across participants (Figure 1A).

### 2.3 Computational modeling and behavioral analyses

To quantify the computational processes underlying exploration–exploitation decisions in dynamic environments and distinguish the effects of cognitive load on different exploration strategies, we compared five reinforcement-learning models: one Q-learning model and four Bayesian learning models based on a Kalman filter (Daw et al., 2006; Wiehler et al., 2021). The Q-learning model updated expected option values based on reward prediction errors and used a softmax function to calculate choice probabilities. Unlike Q-learning, the Kalman filter models tracked both the expected value and uncertainty of each option and adjusted value updating according to current uncertainty. The four Kalman filter models included a basic model, a model with an exploration bonus, a model with a perseveration bonus, and a full model incorporating both bonuses (Supplementary methods 1).

In the Q-learning model, the expected value of each option was updated according to the reward prediction error and transformed into choice probabilities using a softmax function. The parameter β controlled the sensitivity of choices to the estimated option values. Higher β values indicated more consistent value-guided choices, whereas lower values indicated greater choice stochasticity. Unlike Q-learning, the Kalman filter tracked both the expected value and the uncertainty of each option. Uncertainty increased when an option was not sampled and was reduced after the option was selected and feedback was received. Value updating was determined by the prediction error and the trial-wise Kalman gain, allowing learning to depend on the current uncertainty of each option.

To model directed exploration, an exploration bonus parameter, φ, was added in some Kalman filter models. This parameter increased the decision value of an option according to its current uncertainty, such that larger φ values indicated a stronger tendency toward uncertainty-driven directed exploration. Other models additionally included a perseveration parameter, ρ, which captured the tendency to repeat the choice made on the previous trial. In the full model, choice value was determined by expected value, uncertainty bonus, and perseveration bonus and was then transformed into choice probabilities using the softmax function. This framework therefore allowed value sensitivity, directed exploration, and choice perseveration to be modeled separately.

All models were fitted using hierarchical Bayesian estimation in Stan. In Experiment 1, models were compared separately under high and low cognitive load using the Watanabe–Akaike information criterion (WAIC), with lower WAIC values indicating better model performance. Individual parameter estimates from the best-fitting model were then extracted for each load condition, and *t*-tests were used to compare model parameters between high and low cognitive load.

We further evaluated whether the best-fitting model could reproduce participants’ choice behavior. Simulated choices were generated using the fitted model parameters, and behavioral measures from the simulated and observed data were compared. Specifically, stay probabilities were calculated for both observed and simulated choices, and their association was examined. In Experiment 2, model selection was not repeated. Instead, the best-fitting model identified in Experiment 1 was directly applied to the behavioral data from the high- and low-load conditions, and the same parameter comparisons were performed. Model-derived estimates from Experiment 2 were also used for trial classification in the subsequent fMRI analyses.

### 2.4 fMRI preprocessing and GLM analyses

fMRI preprocessing was performed using SPM12 (Friston et al., 1994). Functional images acquired during the first 10 s of each run were discarded to reduce signal instability at the beginning of scanning. The remaining functional images were then corrected for slice timing and head motion using realignment. Each participant’s T1-weighted structural image was coregistered to the functional images and segmented using an East Asian template. The deformation parameters obtained from structural-image segmentation were then used to normalize the functional images to Montreal Neurological Institute (MNI) space, with a voxel size of 3 × 3 × 3 mm^3^ and a bounding box of [−90, −126, −72; 90, 90, 108]. Finally, the normalized images were spatially smoothed using a 6-mm full-width at half-maximum (FWHM) Gaussian kernel. The preprocessed images were used for the subsequent analyses.

We used a hypothesis-driven ROI-based small-volume correction (SVC) approach (Poldrack, 2007; Worsley et al., 1996) to test whether decision-related regions previously identified in exploration–exploitation studies were also involved in directed exploration in the current working-memory-load × decision-making task, and whether their activity was modulated by cognitive load. ROI selection was mainly based on regions consistently reported in previous neuroimaging studies of exploration–exploitation decision-making, including the frontopolar cortex (FP), intraparietal sulcus (IPS), anterior insula (aIns), and dorsal anterior cingulate cortex (dACC). A classic four-armed bandit study first showed that the FP and IPS were involved in exploratory choices (Daw et al., 2006). Later studies further reported increased activity in the aIns and dACC during exploration compared with exploitation (Blanchard & Gershman, 2018), and exploration-related effects in the FP, IPS, aIns, and dACC were also replicated in dynamic four-armed bandit tasks (Chakroun et al., 2020; Wiehler et al., 2021). Wiehler et al. further used 10-mm spherical ROIs centered on previously reported coordinates for SVC analysis, providing a direct methodological reference for the ROI definition used in the present study (Wiehler et al., 2021).

During ROI activation validation, we examined the bilateral homologous regions of these areas to assess the stability of directed-exploration-related activity in the current task. Subsequent connectivity analyses mainly focused on the right-hemisphere decision-related network, including the right frontopolar cortex (rFP), right intraparietal sulcus (rIPS), right anterior insula (raIns), and right dorsal anterior cingulate cortex (rdACC). For the regional load-modulation analysis, we focused on these same four right-hemisphere ROIs to maintain consistency with the subsequent PPI analysis. This analysis was not intended to assess hemispheric lateralization. Because the frontopolar cortex covers a relatively large anatomical area, the search for the FP peak was further restricted to a predefined FP ROI to improve anatomical specificity. The local peak within this restricted ROI was then used for subsequent analyses.

First-level fMRI analyses were performed using the general linear model (GLM) implemented in SPM12. We first constructed GLM1 to examine differences in brain activity across decision types. Trial classification was based on the best-fitting model identified in the behavioral analysis. A trial was classified as an exploitation trial when the chosen option had the highest combination of expected value, uncertainty bonus, and perseveration bonus. All remaining trials were classified as exploration trials. Exploration trials were further divided into directed exploration and random exploration. Trials in which participants selected the option with the highest uncertainty/exploration bonus were classified as DE, whereas the remaining exploration trials were classified as RE. In the subsequent activation analyses, exploitation trials served as the baseline condition for examining brain activity associated with directed exploration relative to exploitation.

To further examine the effect of cognitive load on directed-exploration-related activity, we constructed GLM2 by adding the cognitive-load factor to the decision conditions in GLM1. Trials were modeled separately for the high- and low-load conditions (High Load, HL; Low Load, LL). This model was used to test whether the DE > exploitation effect showed different ROI activation patterns between the two load conditions. At the group level, one-sample *t*-tests were performed by entering each participant’s first-level contrast image into a second-level model to test whether the corresponding contrast differed significantly from zero. The search volume for each ROI was defined as a 10-mm-radius sphere centered on the corresponding peak coordinate.

### 2.5 Psychophysiological interaction analyses

To examine the effect of cognitive load on functional coupling within the decision network related to directed exploration, we further performed psychophysiological interaction (PPI) analyses(Friston et al., 1997). Based on the ROI analyses described above and previous exploration–exploitation studies, four right-hemisphere ROIs were included: the right frontopolar cortex (rFP; 27, 54, 12), right intraparietal sulcus (rIPS; 39, −36, 42), right anterior insula (raIns; 32, 22, −8), and right dorsal anterior cingulate cortex (rdACC; 8, 16, 46). For each ROI, a 5-mm-radius spherical volume of interest (VOI) was centered on the individual local peak, and the corresponding time series was extracted. Each ROI was then used as a seed in a separate PPI model to test task-dependent changes in functional coupling with the remaining ROIs. Two load-related contrasts were examined: an overall decision load contrast (Load_HLgtLL_allDecision), which compared coupling across all decision trials between the high- and low-load conditions, and a directed-exploration-specific contrast (DE_HLgtLL), which compared coupling during high-load DE and low-load DE. Both contrasts were defined as high load > low load; therefore, negative PPI estimates indicated stronger functional coupling under low load. At the group level, one-sample t-tests were performed for the 12 ordered non-self seed–target pairs in each contrast to test whether the PPI contrast estimates differed significantly from zero, with multiple comparisons corrected using the Benjamini–Hochberg false discovery rate (FDR). Candidate connections were further evaluated using permutation tests, bootstrap 95% confidence intervals, and directional Bayes factors as supplementary sensitivity analyses.

### 2.6 Dynamic causal modeling analyses

To further examine directional effective connectivity within the network related to directed exploration, we performed dynamic causal modeling (DCM) based on the PPI results (Friston et al., 2003) Because the PPI analyses showed that load-related changes in functional coupling were mainly observed among the right intraparietal sulcus (rIPS), right anterior insula (raIns), and right dorsal anterior cingulate cortex (rdACC), the subsequent DCM analyses focused on this three-node network.

The DCM analysis was conducted in two steps. The first step aimed to identify the most likely entry point of directed-exploration input into the network. All candidate models included full bidirectional intrinsic connectivity among the rIPS, raIns, and rdACC, together with self-connections for each region. The directed-exploration input was defined as the combined high-load and low-load DE conditions (DE_all = HL-DE + LL-DE). Without specifying a priori which region received the input, candidate models were constructed with DE_all entering the rIPS, raIns, rdACC, or different combinations of these regions. Bayesian model selection was then used to compare model evidence and identify the best-fitting DCM architecture.

In the second step, we used the optimal input structure identified in Step 1 to compare effective connectivity during directed exploration under high and low cognitive load. Separate condition-specific DCMs were constructed for HL-DE and LL-DE, using the same three-node network structure and the same input location. Connection parameters were then extracted for each participant, and differences between the two load conditions were calculated as LL-DE minus HL-DE. For each between-region connection and self-connection, one-tailed tests were used to assess whether the parameter was stronger under low cognitive load. Bayes factors were also calculated as supplementary evidence to evaluate whether effective connectivity related to directed exploration showed an increase under low cognitive load.

## 3. Results

### 3.1 Model comparison and behavioral parameters

We found that the three-parameter model had the lowest WAIC under both high- and low-cognitive-load conditions (Figure 2a). We first examined working memory performance to assess the effectiveness of the cognitive-load manipulation. Participants completed significantly fewer working memory trials correctly under high load than under low load (*t* = 11.5573, *p* < 0.0001; Figure 2d). We then compared the parameters of the best-fitting model between the two load conditions. The parameter phi was significantly higher under low cognitive load than under high cognitive load (*t* = 3.1204, *p* = 0.0044; Figure 2c), indicating that participants were more likely to engage in directed exploration under low cognitive load.

**Figure 2.**
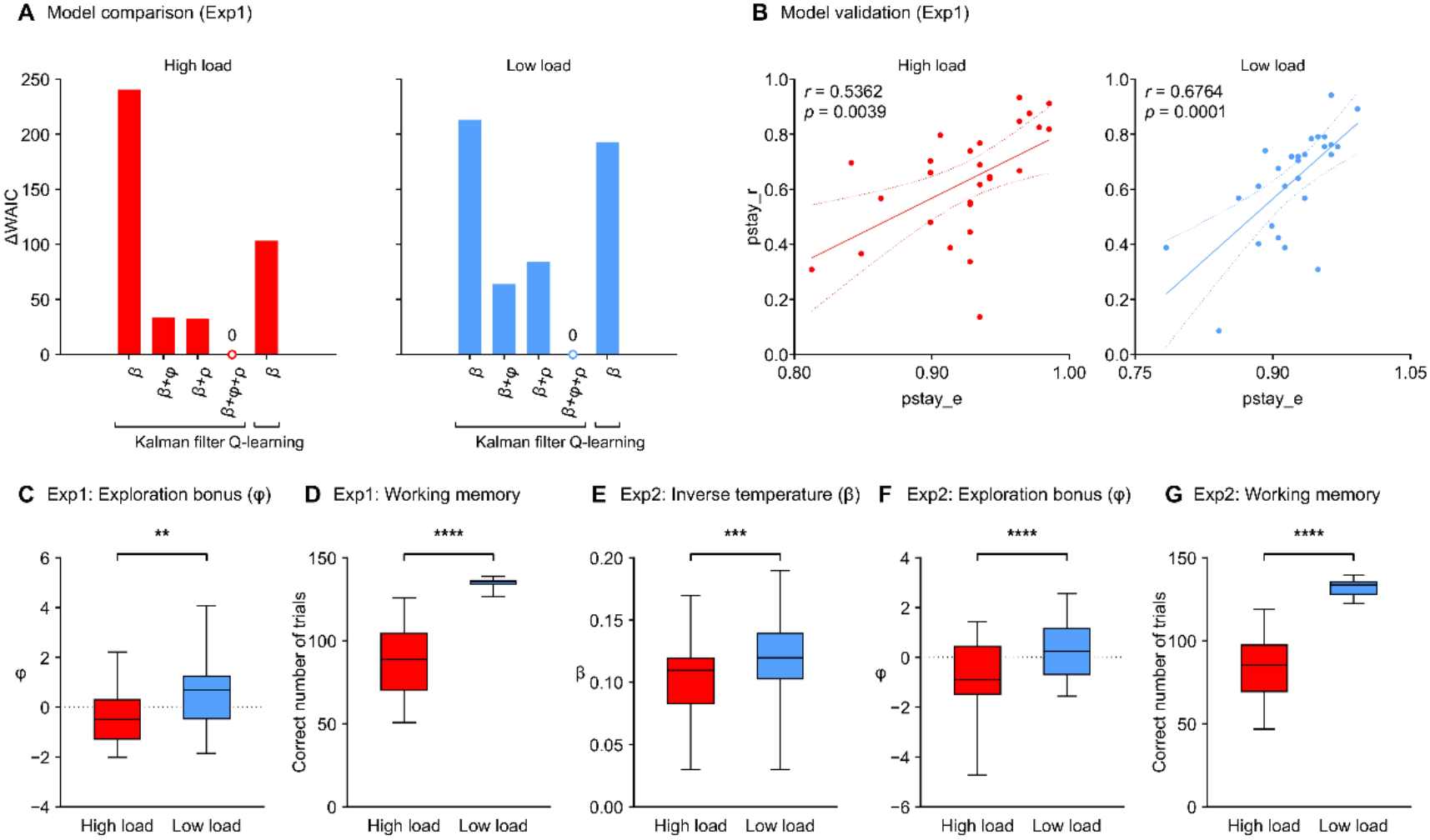
Behavioral model comparison, validation, and parameter results across two experiments. (A) Model comparison in Experiment 1 under high- and low-load conditions based on ΔWAIC. Lower values indicate better model fit. The three-parameter Kalman model showed the best fit under both load conditions. (B) Model validation in Experiment 1. Scatterplots show the relationships between observed (pstay_r) and model-simulated (pstay_e) stay probabilities under high and low load. Significant positive correlations were observed in both conditions. (C) Comparison of the exploration bonus parameter (φ) between load conditions in Experiment 1. φ was significantly higher under low load than under high load. (D) Working memory performance in Experiment 1, showing more correct trials under low load. (E, F) Model parameter comparisons in Experiment 2. Both the inverse temperature parameter and exploration bonus parameter were higher under low load. (G) Working memory performance in Experiment 2, showing more correct trials under low load. Red and blue indicate high- and low-load conditions, respectively. \*\**p* < 0.01, \*\*\**p* < 0.001, \*\*\*\**p* < 0.0001. Abbreviations: WAIC, Watanabe–Akaike information criterion; φ, exploration bonus parameter; pstay_r, observed stay probability; pstay_e, expected stay probability predicted by the model.

We further performed parameter recovery for the best-fitting model and calculated the correlation between the observed pstay and the pstay generated from the simulated data. The observed pstay was defined as the probability of choosing the same option as in the previous trial. Significant positive correlations were found under both high cognitive load (*r* = 0.5362, *p* = 0.0039) and low cognitive load (*r* = 0.6764, *p* = 0.0001; Figure 2b).

In Experiment 2, we directly applied the best-fitting model identified in Experiment 1. Consistent with Experiment 1, working memory performance was significantly lower under high load than under low load (*t* = 18.2590, *p* < 0.0001; Figure 2g), further supporting the effectiveness of the cognitive-load manipulation. We again compared model parameters between the two cognitive-load conditions. Consistent with Experiment 1, phi was significantly higher under low cognitive load than under high cognitive load (*t* = 5.8250, *p* < 0.0001, Figure 2f). We also found that beta was significantly higher under low cognitive load (*t* = 4.2104, *p* = 0.0001, Figure 2e).

### 3.2 GLM results

In GLM1, we examined the main effect of DE > exploitation. ROI-based small-volume correction showed significant activation in the predefined FP, IPS, aIns, and dACC ROIs for the DE > exploitation contrast. These results indicate that these classic decision-related regions remained involved in directed exploration in the current cognitive-load × decision-making task (Table 1; Figure 3a).

**Table 1.** ROI analysis.

| **Reference** | **Location** | **Coordinates** | **MainEffect** | **Highload** | **Lowload** |
|---|---|---|---|---|---|
|  |  | **(x/y/z)** | **DE>exploit** | **DE>exploit** | **DE>exploit** |
| Present study | R FP | 27, 54, 12 | *p* = 0.001 | *p* = 0.026 | *p* = 0.002 |
| Present study | L FP | -36, 51, 6 | *p* < 0.001 | *p* = 0.003 | *p* < 0.001 |
| Daw et al. (2006) | R IPS | 39, -36, 42 | *p* < 0.001 | *p* = 0.018 | *p* = 0.001 |
| Daw et al. (2006) | L IPS | -29, -33, 45 | *p* < 0.001 | No cluster | *p* < 0.001 |
| Blanchard and Gershman (2018) | R aIns | 32, 22, -8 | *p* < 0.001 | *p* = 0.019 | *p* < 0.001 |
| Blanchard and Gershman (2018) | L aIns | -30, 16, -8 | *p* < 0.001 | *p* = 0.041 | *p* = 0.002 |
| Blanchard and Gershman (2018) | R dACC | 8, 16, 46 | *p* < 0.001 | *p* = 0.002 | *p* < 0.001 |

**Figure 3.**
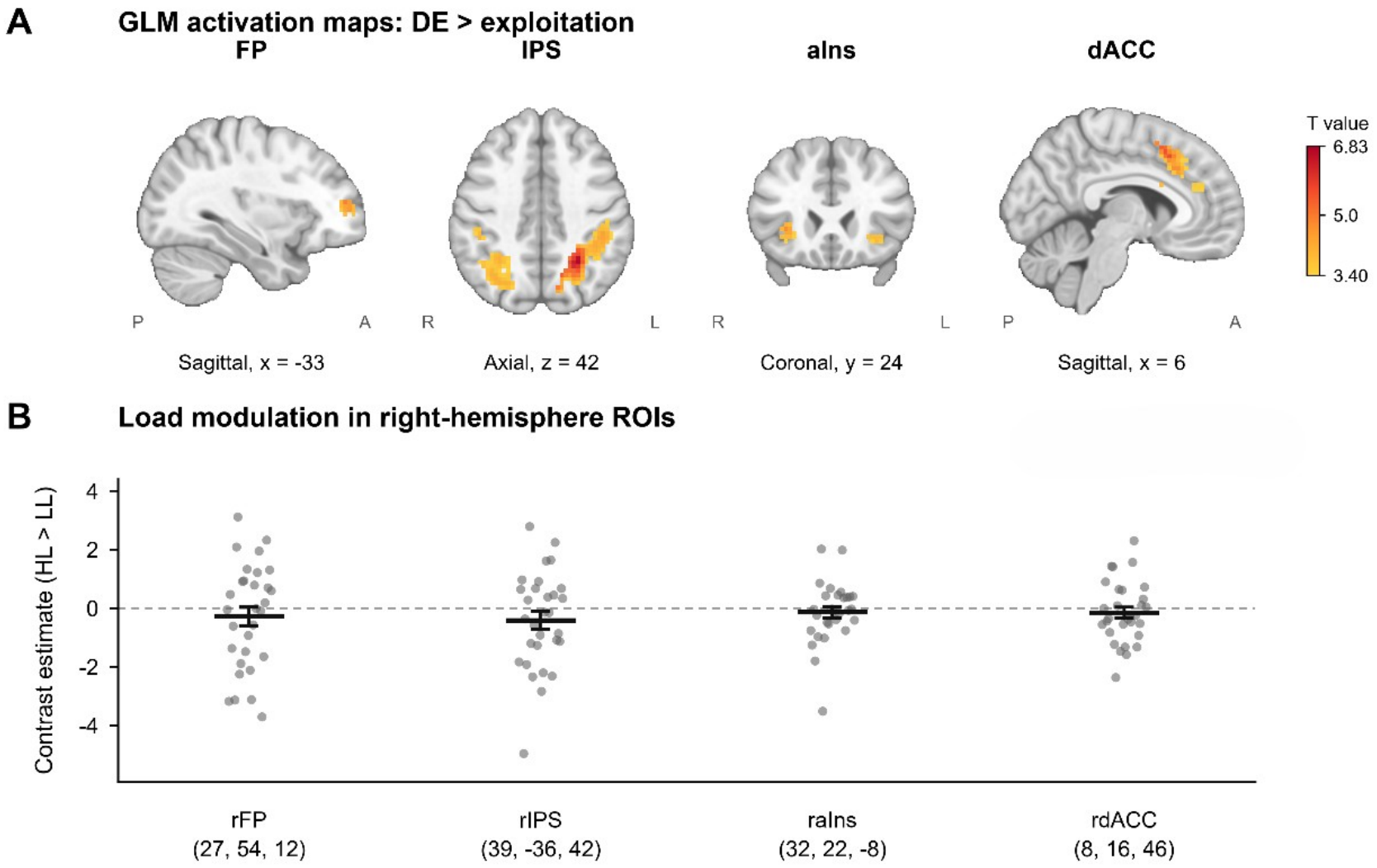
ROI activation during directed exploration and its modulation by cognitive load. (A) ROI-masked activation maps for the DE > exploitation contrast in the frontopolar cortex (FP), intraparietal sulcus (IPS), anterior insula (aIns), and dorsal anterior cingulate cortex (dACC). Color indicates *t* values. (B) Load modulation of directed-exploration-related activation in the right-hemisphere ROIs. Contrast estimates were extracted from the high-load > low-load contrast of the DE > exploitation effect. Gray dots represent individual participants, and black lines with error bars represent the group mean and its uncertainty. None of the four ROIs showed a significant load-modulation effect; rIPS showed a nonsignificant trend toward stronger activation under low cognitive load (*p* = 0.089). Abbreviations: DE, directed exploration; rFP, right frontopolar cortex; rIPS, right intraparietal sulcus; raIns, right anterior insula; rdACC, right dorsal anterior cingulate cortex; HL, high load; LL, low load.

To further test whether directed-exploration-related activity was modulated by cognitive load, we constructed 10-mm-radius spherical ROIs around the peak coordinates of the key right-hemisphere regions and extracted the mean contrast estimate for each participant from the high-load > low-load contrast of the DE > exploitation effect. The peak coordinates were rFP [27, 54, 12], rIPS [39, −36, 42], raIns [32, 22, −8], and rdACC [8, 16, 46]. None of the four right-hemisphere ROIs showed a load-modulation contrast that differed significantly from zero. A one-tailed test showed a trend toward a stronger DE > exploitation effect in the rIPS under low load than under high load, but this effect did not reach statistical significance (*p* = 0.089; Figure 3b).

### 3.3 PPI analysis

The PPI analysis first examined the effect of cognitive load during the overall decision-making process. In the overall decision load contrast, the connection from raIns as the seed to rIPS as the target showed a negative PPI estimate (mean = −0.5515, *t* = −2.3599, two-sided *p* = 0.0252, Cohen’s d = −0.4309; Figure 4a, PPI1), indicating stronger functional coupling under low cognitive load. Sensitivity analyses for this connection showed a two-sided permutation *p* = 0.0250, a bootstrap 95% CI of [−1.0130, −0.1030], and an approximate directional Bayes factor of 5.03 (Figure 4b, PPI1). The connection from rdACC as the seed to rIPS as the target also showed a negative effect (mean = −0.3124, *t* = −2.2403, two-sided *p* = 0.0329, Cohen’s d = −0.4090; Figure 4a, PPI1), with a two-sided permutation *p* = 0.0330, a bootstrap 95% CI of [−0.5790, −0.0460], and an approximate directional Bayes factor of 3.94 (Figure 4b, PPI1).

**Figure 4.**
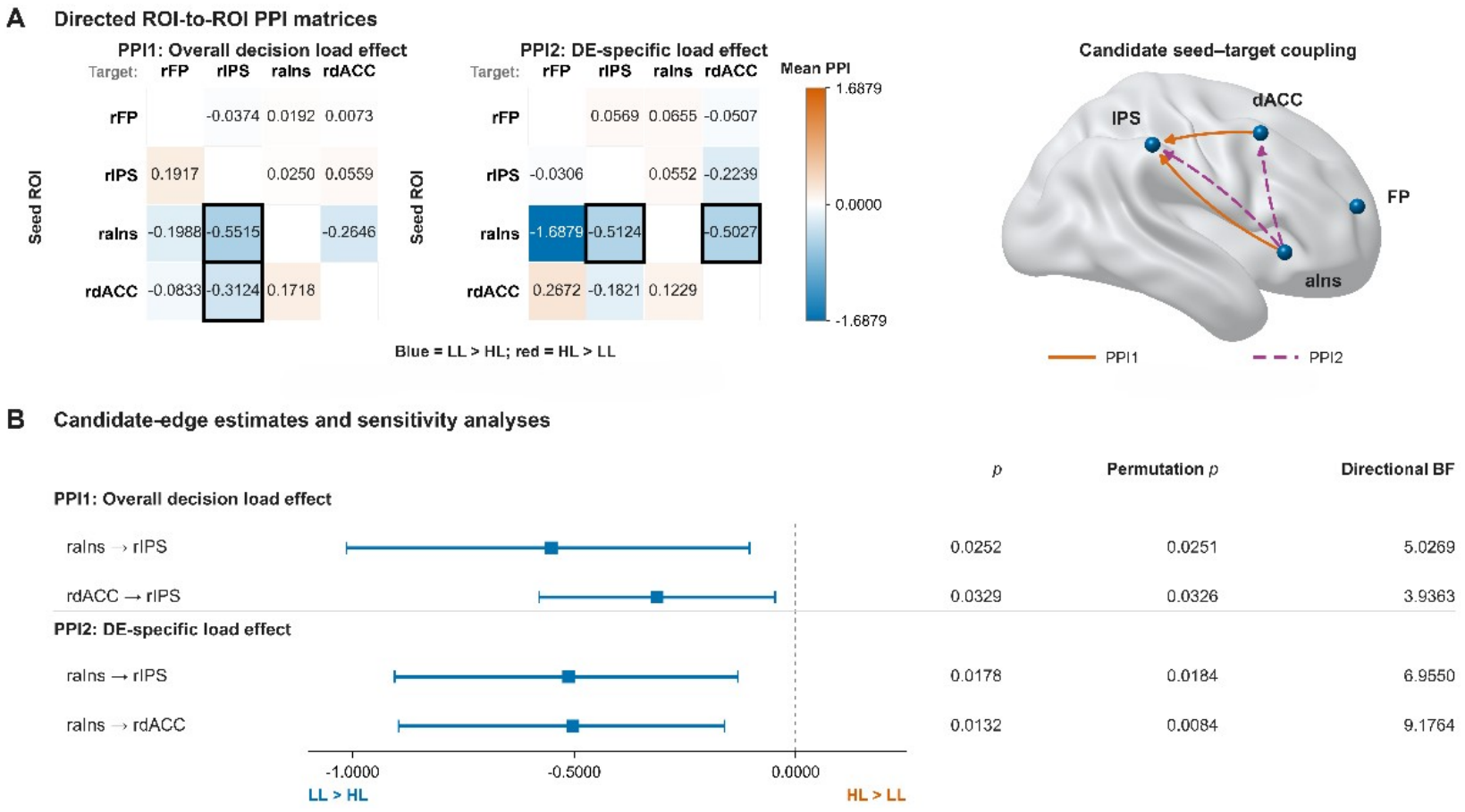
Load-related changes in functional coupling from the PPI analyses. (A) Directed ROI-to-ROI PPI matrices for the overall decision load effect (PPI1) and the directed-exploration-specific load effect (PPI2). Matrix values represent PPI contrast estimates for each seed–target connection. Blue indicates stronger coupling under low load than high load, whereas red indicates stronger coupling under high load than low load. Black outlines indicate candidate connections with uncorrected *p* < 0.05. The brain network on the right shows these candidate connections on a lateral brain surface. Orange solid lines represent PPI1, and purple dashed lines represent PPI2. Arrows indicate seed–target ordering rather than causal direction. (B) Candidate-edge estimates and sensitivity analyses for the significant connections identified in Panel A. Points and horizontal bars show the mean PPI contrast estimate and its 95% confidence interval. Negative estimates indicate stronger coupling under low load. The corresponding two-sided parametric p values, permutation p values, and directional Bayes factors are shown on the right. Abbreviations: PPI, psychophysiological interaction; PPI1, overall decision load effect; PPI2, directed-exploration-specific load effect; BF, Bayes factor.

For the load comparison during directed exploration, that is, the DE_HLgtLL contrast, the connection from raIns to rIPS showed a negative effect (mean = −0.5124, *t* = −2.5133, two-sided *p* = 0.0178, Cohen’s d = −0.4589; Figure 4a, PPI2). Sensitivity analyses showed a two-sided permutation *p* = 0.0180, a bootstrap 95% CI of [−0.9050, −0.1300], and an approximate directional Bayes factor of 6.95 (Figure 4b, PPI2). The connection from raIns to rdACC also showed a negative effect (mean = −0.5027, *t* = −2.6400, uncorrected two-sided parametric *p* = 0.0132, Cohen’s d = −0.4820; Figure 4a, PPI2), with a two-sided permutation *p* = 0.0080, a bootstrap 95% CI of [−0.8960, −0.1610], and an approximate directional Bayes factor of 9.18 (Figure 4b, PPI2).

### 3.4 DCM analysis

To identify the most likely entry point of directed-exploration-related input within the rIPS–raIns– rdACC network, we compared candidate DCMs with different DE input locations (Figure 5a). Bayesian model selection favored the model with DE input entering the rIPS, with a fixed-effects posterior probability of approximately 1.00 and a random-effects exceedance probability of 0.724 (Figure 5b).

**Figure 5.**
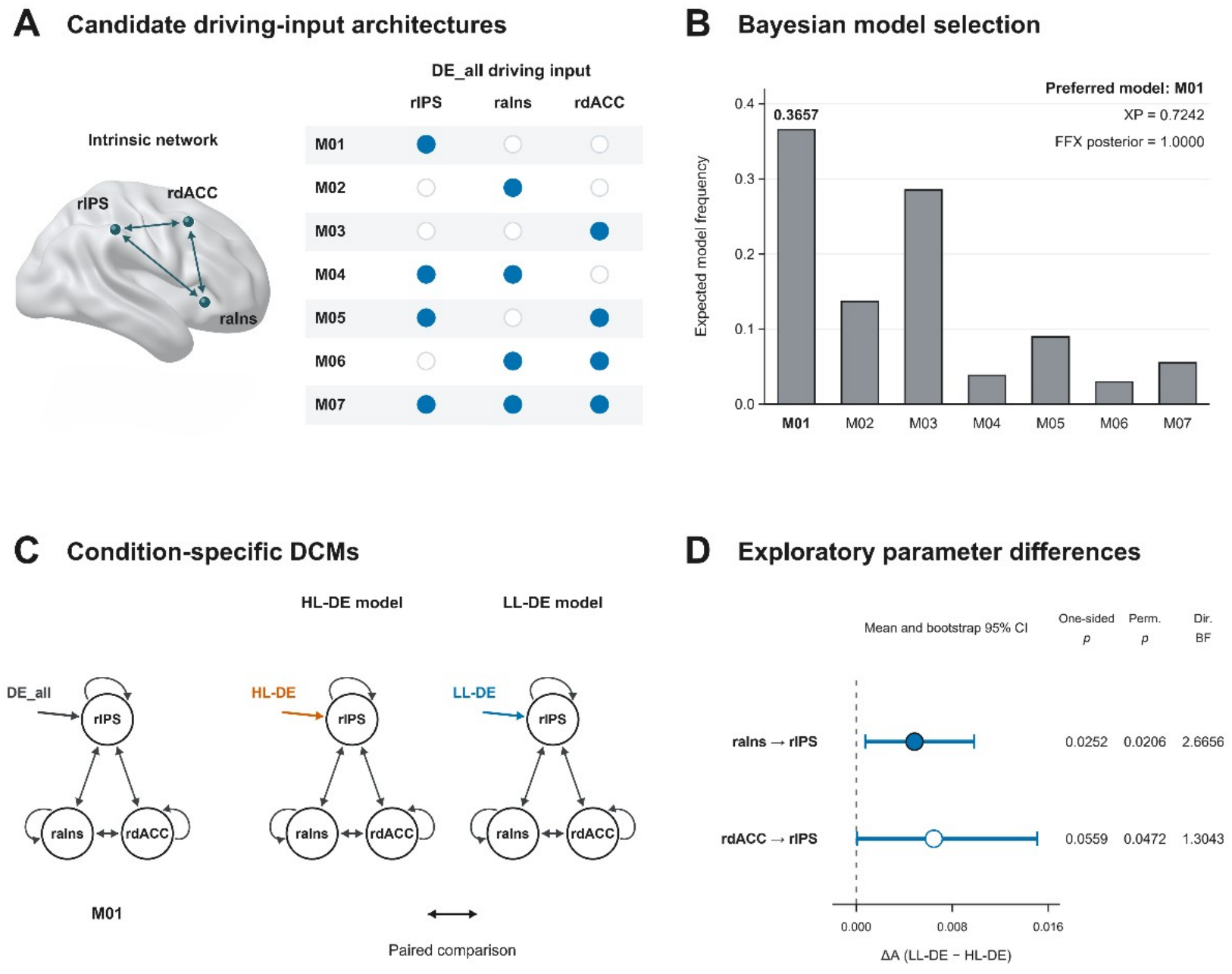
DCM results for the directed-exploration-related network. (A) Candidate driving-input architectures tested in the three-node network comprising the rIPS, raIns, and rdACC. All models included full bidirectional intrinsic connections and self-connections, while differing in the location of the DE_all driving input. (B) Bayesian model selection results for the candidate models. Model M01, in which DE_all entered the rIPS, showed the highest model evidence in both fixed-effects and random-effects analyses. (C) Condition-specific DCMs used for parameter comparison. After selecting M01 as the optimal architecture, separate models were constructed for high-load directed exploration (HL-DE) and low-load directed exploration (LL-DE). Parameter differences were then calculated as LL-DE minus HL-DE. (D) Parameter differences for candidate connections showing stronger effective connectivity under low load. Points and horizontal bars represent the mean difference and bootstrap 95% confidence interval. Positive values indicate stronger effective connectivity in the LL-DE model than in the HL-DE model. One-sided p values, permutation p values, and directional Bayes factors are shown on the right. Abbreviations: DE_all, combined directed exploration input across load conditions; FFX, fixed effects; XP, exceedance probability.

After identifying DE input → rIPS as the optimal input structure, we further constructed separate condition-specific DCMs for high-load DE and low-load DE and compared their effective connectivity parameters. Specifically, for each participant, parameter differences were calculated as low-load DE minus high-load DE (LL-DE minus HL-DE; Figure 5c). Based on one-tailed tests together with Bayes factor evidence, several connections showed a tendency toward stronger parameters under low cognitive load. The raIns → rIPS connection was stronger in the low-load DE model than in the high-load DE model (*t* = 2.041, Cohen’s d = 0.373). The directional permutation test was significant (*p* = 0.021), and the Bayes factor was greater than 1 (BF = 2.67), suggesting that this connection may be enhanced under low cognitive load. The rdACC → rIPS connection showed a similar trend (*t* = 1.640, Cohen’s d = 0.299, permutation test *p* = 0.047, BF = 1.30; Figure 5d), suggesting that effective connectivity from the rdACC to the rIPS may also be stronger under low cognitive load. Other between-region connections and self-connections did not show stable enhancement under low cognitive load.

## 4. Discussion

This study combined computational modeling, task-based fMRI, PPI, and DCM to examine how cognitive load affects exploration strategies and related brain network interactions in dynamic environments. Across two experiments, higher cognitive load reduced uncertainty-driven DE, while Experiment 2 further showed increased choice stochasticity under high load, suggesting that greater cognitive demands may shift behavior from goal-directed information seeking toward more random exploration. Neuroimaging results showed that although DE, relative to exploitation, engaged the FP, IPS, aIns, and dACC, these regions showed no clear load-related differences in regional activation. In contrast, PPI suggested that cognitive load modulated functional coupling among exploration-related regions, particularly between the aIns and IPS. DCM further supported the IPS as a candidate recipient of DE-related driving input and suggested higher aIns-to-IPS effective-connectivity estimates under low load. Together, these findings suggest that reduced DE and increased choice stochasticity under cognitive load may be associated with selective reconfiguration of interactions within an IPS-related decision network.

First, the behavioral and computational modeling results support separable effects of cognitive resources on directed and random exploration. Directed exploration requires individuals to track uncertainty and assign additional information value to uncertain options, whereas random exploration more closely reflects reduced value-guided choice or increased choice variability (Wilson et al., 2014). Consistent with this framework, participants showed a higher directed-exploration parameter under low cognitive load, suggesting more effective use of uncertainty when cognitive resources were available. Across both experiments, higher cognitive load consistently reduced the exploration bonus, whereas increased choice stochasticity, reflected by lower β, was observed only in Experiment 2. Similar effects have been reported previously: higher cognitive load reduced directed exploration and increased random exploration, partly through impaired integration of reward and information value (Cogliati Dezza et al., 2019), while increased memory demands reduced strategic exploration of unsampled options and increased reliance on known information (Brown et al., 2022). Together, these findings suggest that cognitive load does not simply change the general tendency to explore, but affects the ability to translate uncertainty into goal-directed exploration. This interpretation is also consistent with recent work on individual differences in working memory capacity and exploration strategies. Witte et al. found that model-derived exploration measures were associated with working memory capacity, suggesting that part of the variability in exploration parameters may reflect differences in cognitive resources (Witte et al., 2025). Notably, exploration parameters showed only low-to-moderate consistency across different bandit tasks, indicating that they should not be viewed as fixed individual traits. By experimentally manipulating cognitive load within the same individuals, the present study provides more direct evidence that available cognitive resources can alter the computational processes underlying exploration strategies.

At the neural level, the present study first replicated the finding that directed exploration, relative to exploitation, involves regions including the FP, IPS, aIns, and dACC. This is consistent with the frontoparietal exploration system identified in the classic four-armed bandit stud (Daw et al., 2006), as well as later studies that repeatedly observed exploration-related effects in the FP, IPS, aIns, and dACC during dynamic bandit tasks (Chakroun et al., 2020; Wiehler et al., 2021). Using a model-driven trial classification approach highly similar to that used in the present study, Wiehler et al. likewise found that directed exploration > exploitation reliably activated the FP, IPS, aIns, and dACC. Therefore, the present study shows that this classic exploration-related neural system can still be reliably identified in a dual-task context with additional cognitive load, indicating that the directed exploration captured by the current paradigm has a neural basis consistent with previous studies. However, an important finding was that although directed exploration differed clearly between high- and low-cognitive-load conditions, directed-exploration-related univariate activation did not show an equally clear load effect. This incomplete correspondence between behavior and regional activation does not imply that cognitive load had no effect on the exploration system, because the modulation of complex decision-making may also depend on information exchange between regions. Wiehler et al. observed a similar pattern in gambling disorder: despite clear group differences in directed exploration, several classic exploration-related ROIs did not show stable univariate group effects, whereas DCM connectivity parameters provided additional group information (Wiehler et al., 2021). Recent work has also shown that increasing computational demands can alter both regional activity and connectivity among the aIns, dACC, and parietal cortex (Franco et al., 2024). Therefore, the combination of relatively stable regional activation and altered connectivity patterns in the present study is more consistent with a network reconfiguration account. Cognitive load may primarily change how exploration-related regions coordinate information, rather than simply changing whether these regions are involved in exploration.

The PPI results further support this interpretation. Load-related changes in functional coupling were mainly observed within the decision network involving the IPS, aIns, and dACC. This is consistent with recent evidence showing that cognitive computational demands jointly modulate both regional activity and functional connectivity in the aIns, dACC, and parietal cortex (Franco et al., 2024). A recent fMRI meta-analysis of decision uncertainty, including 76 studies and 4,186 participants, further identified the anterior insula and parietal cortex as among the most consistently engaged regions across tasks, while the ACC/aIns system was repeatedly involved in integrating uncertainty information with control demands (Timashkov et al., 2025). Beyond the exploration literature, increased cognitive effort has also been shown to recruit the dACC, aIns, and IPS and to alter functional connectivity between these regions (Aben et al., 2020). Therefore, the connectivity findings in the present study may indicate that higher cognitive load disrupts the coordination between maintaining and comparing uncertainty information and adjusting behavior according to current demands, thereby shifting choice from more purposeful directed exploration toward more random behavior. It should be emphasized, however, that PPI reflects task-dependent statistical coupling rather than directional causal influence. Therefore, interpretation of the direction of information flow requires further evidence from the DCM results.

DCM further revealed how information is organized within the directed-exploration-related network and how this organization is modulated by cognitive load. First, among the candidate input models, the model in which the IPS received the driving input received the strongest support, consistent with the findings of Wiehler et al. in a four-armed restless bandit task (Wiehler et al., 2021). Together with previous evidence linking the parietal cortex to uncertainty representation, evidence integration, and dynamic decision-making (Leach et al., 2025; Valdebenito-Oyarzo et al., 2024), this finding further suggests that the IPS may serve as an important candidate node through which uncertainty-related information enters the exploration process. Furthermore, the second-stage DCM suggested that cognitive load may selectively modulate effective connectivity within this network. Compared with high cognitive load, the raIns → rIPS and rdACC → rIPS connections were stronger under low cognitive load, in the same direction as the stronger directed exploration observed behaviorally. This suggests that when more cognitive resources are available, signals from salience- and control-related regions may more effectively influence uncertainty- and evidence-related processing in the IPS, thereby supporting the transformation of uncertainty into goal-directed exploration. Importantly, these load effects were not observed across all connections, but were selective to specific pathways. This pattern suggests that cognitive load may reconfigure information exchange within the exploration network rather than uniformly enhancing or suppressing the entire system. Such selective modulation is consistent with task-demand-dependent connectivity changes reported in cognitive control research, where changes in cognitive resource demands can influence behavior by reorganizing communication between regions rather than by producing parallel changes in activity across all nodes (Aben et al., 2020). Together with the load-related functional coupling observed in the PPI analysis, these findings support the interpretation that lower cognitive load may provide a more favorable network state for directed exploration, allowing uncertainty, behavioral salience, and control demands to be integrated more effectively within the IPS–aIns–dACC decision system. As cognitive load increases, this coordination may weaken and be accompanied by reduced directed exploration.

From a broader perspective, the mechanism identified in the present study—limited cognitive resources, reduced directed exploration, increased random exploration, and altered network coordination—may provide a useful basis for understanding abnormalities in adaptive decision-making across psychiatric disorders. Imbalance in the exploration–exploitation trade-off is not specific to a single disorder. Individuals with gambling disorder show reduced directed exploration, and connectivity patterns within the exploration network can distinguish patients from healthy individuals (Wiehler et al., 2021). Patients with schizophrenia, in contrast, may show increased random exploration, which has been associated with disorganization symptoms and inflammatory markers (Cathomas et al., 2021). A recent study using a volatile four-armed bandit task also found that individuals with remitted major depressive disorder showed reduced punishment learning and choice stochasticity, highlighting differences in learning and decision-making processes across clinical states (Moses-Payne et al., 2026). A previous study further showed that abnormal stay–switch behavior under uncertain feedback was influenced by feedback sensitivity and serotonergic medication status, and was also related to clinical severity (Apergis-Schoute et al., 2024) A systematic review published in 2025 similarly concluded that psychiatric disorders do not show a uniform pattern of either “increased exploration” or “reduced exploration.” Instead, abnormalities may occur in different computational components, including random exploration, goal-directed exploration, feedback learning, and behavioral persistence (Jami et al., 2025).

In this context, the value of the present study is not to equate high cognitive load in healthy individuals with psychiatric disorders, but rather to provide an experimentally controllable model of a basic mechanism. Specifically, when the cognitive resources available for decision-making are temporarily reduced, even healthy individuals may show a behavioral shift from directed exploration toward more random exploration (Cogliati Dezza et al., 2019). This raises the possibility that exploration abnormalities in some clinical populations may partly arise from changes in cognitive resources, information integration, or network coordination, rather than being fully explained by altered reward sensitivity itself (Cathomas et al., 2021; Wiehler et al., 2021).

This perspective is also consistent with the current computational psychiatry framework emphasizing transdiagnostic computational processes (Huys et al., 2016). Similar behavioral abnormalities may result from different underlying mechanisms, while impairment in the same computational function may cut across traditional diagnostic boundaries. Therefore, future studies examining dynamic changes in exploration parameters and network connectivity across different levels of cognitive load in clinical populations may help distinguish relatively stable disorder-related characteristics from resource-dependent abnormalities that vary with cognitive state.

## 5. Limitations

Several limitations should be considered. First, the present study used a hypothesis-driven ROI approach. Although this increased sensitivity for testing previously established hypotheses about the exploration-related network, it limited the identification of other potentially relevant brain regions and whole-brain networks. Therefore, the current conclusions should be mainly restricted to the predefined decision-related system. Second, the dynamic four-armed bandit task was well suited to our aim of examining exploration in continuously changing reward environments.

However, unlike the Horizon Task, it does not explicitly manipulate information availability and future exploration opportunities to distinguish exploration strategies. Therefore, the exploration bonus parameter φ should be interpreted as a model-derived measure of uncertainty-guided exploration. Future studies could combine Horizon Task manipulations with dynamic reward environments to better distinguish directed and random exploration. Thirdly, although several PPI connections showed consistent effects across parametric tests, permutation tests, bootstrap analyses, and Bayes factors, none survived FDR correction. These complementary analyses provide converging evidence for candidate connections but do not fully address the multiple-comparison issue. Furthermore, although DCM model selection favored the IPS as the input node, the evidence for load-related differences in effective connectivity was relatively weak (BF < 3).

Thus, these findings suggest potentially meaningful changes in IPS-centered network interactions, which warrant further validation in independent samples. Fourthly, 10 of the 40 participants in Experiment 2 had no trials classified as directed exploration and were excluded from the fMRI analyses. Although the main behavioral effects remained significant in the retained sample (Supplementary Results 1), this does not eliminate potential selection bias. The imaging sample may therefore overrepresent individuals showing model-classified directed exploration, limiting the generalizability of the neural findings. Future studies could use task designs or analysis approaches that accommodate a broader range of exploration behavior. Finally, the present study was based on an acute cognitive-load manipulation in healthy individuals and therefore cannot directly address long-term neural adaptations in psychiatric disorders. Future studies combining clinical samples, individual differences in working memory capacity, pharmacological manipulations, and longitudinal or multi-time-point designs may help clarify the causal relationships among cognitive resources, neurotransmitter systems, and exploration-related networks.

## 6. Conclusion

The present study shows that cognitive load alters the exploration–exploitation trade-off: higher load consistently reduced model-derived directed exploration across both experiments. Neuroimaging results further suggest that this shift is more closely related to changes in interactions within the exploration-related network than to simple changes in regional activation. PPI and DCM findings together support an important role of the IPS–aIns–dACC network, with the IPS serving as a candidate input node for directed-exploration-related information. Overall, these findings link cognitive resources, exploration strategies, and network organization, and provide a mechanistic basis for understanding exploration–exploitation abnormalities in psychiatric disorders.

## CRediT authorship contribution statement

**Haowei Dai:** Writing – review & editing, Writing – original draft, Visualization, Software, Methodology, Formal analysis, Data curation, Conceptualization. **Linhui Wang:** Investigation, Data curation. **Xueping Yin:** Investigation. **Xiangyi Liang:** Investigation. **Yutong Ying:** Investigation. **Yuanyuan Zeng:** Investigation. **Lijing Niu:** Investigation. **Jonathan P. Roiser:** Conceptualization, Supervision. **Xiangliang Tan:** Methodology, Resources. **Ruibin Zhang:** Conceptualization, Methodology, Funding acquisition, Project administration, Supervision, Writing – review & editing.

## Data availability

The datasets used and/or analyzed during the present study are available from the corresponding author upon reasonable request.

## References

Aben, B., Calderon, C. B., Bussche, E. V. den, & Verguts, T. (2020). Cognitive Effort Modulates Connectivity between Dorsal Anterior Cingulate Cortex and Task-Relevant Cortical Areas. Journal of Neuroscience, 40(19), 3838~3848. 10.1523/JNEUROSCI.2948-19.2020

Apergis-Schoute, A. M., van der Flier, F. E., Ip, S. H. Y., Kanen, J. W., Vaghi, M. M., Fineberg, N. A., Sahakian, B. J., Cardinal, R. N., & Robbins, T. W. (2024). Perseveration and Shifting in Obsessive-Compulsive Disorder as a Function of Uncertainty, Punishment, and Serotonergic Medication. Biological Psychiatry Global Open Science, 4(1), 326~335. 10.1016/j.bpsgos.2023.06.004

Blanchard, T. C., & Gershman, S. J. (2018). Pure correlates of exploration and exploitation in the human brain. Cognitive, Affective, & Behavioral Neuroscience, 18(1), 117~126. 10.3758/s13415-017-0556-2

Bond, K., Dunovan, K., Porter, A., Rubin, J. E., & Verstynen, T. (2021). Dynamic decision policy reconfiguration under outcome uncertainty. eLife, 10, e65540. 10.7554/eLife.65540

Brown, V. M., Hallquist, M. N., Frank, M. J., & Dombrovski, A. Y. (2022). Humans adaptively resolve the explore-exploit dilemma under cognitive constraints: Evidence from a multi-armed bandit task. Cognition, 229, 105233. 10.1016/j.cognition.2022.105233

Bullmore, E., & Sporns, O. (2009). Complex brain networks: Graph theoretical analysis of structural and functional systems. Nature Reviews Neuroscience, 10(3), 186~198. 10.1038/nrn2575

Cai, W., & Menon, V. (2025). Heterogeneity of human insular cortex: Five principles of functional organization across multiple cognitive domains. Brain structure & function, 230(8), 161. 10.1007/s00429-025-03027-y

Cathomas, F., Klaus, F., Guetter, K., Chung, H.-K., Raja Beharelle, A., Spiller, T. R., Schlegel, R., Seifritz, E., Hartmann-Riemer, M. N., Tobler, P. N., & Kaiser, S. (2021). Increased random exploration in schizophrenia is associated with inflammation. Npj Schizophrenia, 7(1), 6. 10.1038/s41537-020-00133-0

Chakroun, K., Mathar, D., Wiehler, A., Ganzer, F., & Peters, J. (2020). Dopaminergic modulation of the exploration/exploitation trade-off in human decision-making. eLife, 9, e51260. 10.7554/eLife.51260

Chan, V. K. S., Wong, N. H. L., Woo, T.-F., Watanabe, K., Haruno, M., Law, C.-K., & Chau, B. K. H. (2026). Distinct contributions of prefrontal, parietal, and cingulate signals to exploratory decisions. Communications Biology, 9(1), 272. 10.1038/s42003-026-09550-w

Cogliati Dezza, I., Cleeremans, A., & Alexander, W. (2019). Should we control? The interplay between cognitive control and information integration in the resolution of the exploration-exploitation dilemma. Journal of Experimental Psychology: General, 148(6), 977~993. 10.1037/xge0000546

Cohen, J. D., McClure, S. M., & Yu, A. J. (2007). Should I stay or should I go? How the human brain manages the trade-off between exploitation and exploration. Philosophical Transactions of the Royal Society B: Biological Sciences, 362(1481), 933~942. 10.1098/rstb.2007.2098

Cole, M. W., Reynolds, J. R., Power, J. D., Repovs, G., Anticevic, A., & Braver, T. S. (2013). Multi-task connectivity reveals flexible hubs for adaptive task control. Nature neuroscience, 16(9), 1348~1355. 10.1038/nn.3470

Crawford, J. L., Brough, R. E., Eisenstein, S. A., Peelle, J. E., & Braver, T. S. (2024). Generalized Encoding of the Relative Subjective Value of Cognitive Effort in the Dorsal ACC. The Journal of Neuroscience, 44(38), e0367242024. 10.1523/JNEUROSCI.0367-24.2024

Daw, N. D., O’Doherty, J. P., Dayan, P., Seymour, B., & Dolan, R. J. (2006). Cortical substrates for exploratory decisions in humans. Nature, 441(7095), 876~879. 10.1038/nature04766

Duan, D., Wang, Y., Dai, H., & Zhang, R. (2026). Working memory capacity is linked to directed exploration: Behavioral, computational, and electrophysiological evidence (页 2026.11.08.765180). bioRxiv. 10.64898/2026.11.08.765180

Duncan, J. (2010). The multiple-demand (MD) system of the primate brain: Mental programs for intelligent behaviour. Trends in Cognitive Sciences, 14(4), 172~179. 10.1016/j.tics.2010.01.004

Franco, J. P., Bossaerts, P., & Murawski, C. (2024). The neural dynamics associated with computational complexity. PLOS Computational Biology, 20(9), e1012447. 10.1371/journal.pcbi.1012447

Friston, K. J., Buechel, C., Fink, G. R., Morris, J., Rolls, E., & Dolan, R. J. (1997). Psychophysiological and modulatory interactions in neuroimaging. NeuroImage, 6(3), 218~229. 10.1006/nimg.1997.0291

Friston, K. J., Harrison, L., & Penny, W. (2003). Dynamic causal modelling. NeuroImage, 19(4), 1273~1302. 10.1016/s1053-8119(03)00202-7

Friston, K. J., Holmes, A. P., Worsley, K. J., Poline, J.-P., Frith, C. D., & Frackowiak, R. S. J. (1994). Statistical parametric maps in functional imaging: A general linear approach. Human Brain Mapping, 2(4), 189~210. 10.1002/hbm.460020402

Gershman, S. J. (2018). Deconstructing the human algorithms for exploration. Cognition, 173, 34 42. 10.1016/j.cognition.2017.12.014

Harms, M. B., Xu, Y., Green, C. S., Woodard, K., Wilson, R., & Pollak, S. D. (2024). The Structure and Development of Explore-Exploit Decision Making. Cognitive psychology, 150, 101650. 10.1016/j.cogpsych.2024.101650

Huettel, S. A., Song, A. W., & McCarthy, G. (2005). Decisions under Uncertainty: Probabilistic Context Influences Activation of Prefrontal and Parietal Cortices. Journal of Neuroscience, 25(13), 3304~3311. 10.1523/JNEUROSCI.5070-04.2005

Huys, Q. J. M., Maia, T. V., & Frank, M. J. (2016). Computational psychiatry as a bridge from neuroscience to clinical applications. Nature Neuroscience, 19(3), 404~413. 10.1038/nn.4238

Jami, A., Abbaszade, S., & Vahabie, A.-H. (2025). A review on exploration–exploitation trade-off in psychiatric disorders. BMC Psychiatry, 25(1), 420. 10.1186/s12888-025-06837-w

Kolling, N., Behrens, T. E. J., Mars, R. B., & Rushworth, M. F. S. (2012). Neural Mechanisms of Foraging. Science, 336(6077), 95~98. 10.1126/science.1216930

Leach, S. C., Hollow, H., Jiang, J., & Hwang, K. (2025). Frontoparietal Hubs Leverage Probabilistic Representations and Integrated Uncertainty to Guide Cognitive Flexibility. Journal of Neuroscience, 45(49). 10.1523/JNEUROSCI.0989-25.2025

Lloyd, A., McKay, R., & Furl, N. (2025). Stochastic decisions support optimal foraging of volatile environments, and are disrupted by anxiety. Cognitive, Affective, & Behavioral Neuroscience, 25(3), 868~885. 10.3758/s13415-024-01256-y

Menon, V., & Uddin, L. Q. (2010). Saliency, switching, attention and control: A network model of insula function. Brain structure & function, 214(5~6), 655~667. 10.1007/s00429-010-0262-0

Moses-Payne, M., Zhang, R., Mkrtchian, A., Valton, V., Gray, A., Kieslich, K., VanUrk, S., Samborska, V., Halahakoon, D. C., Dayan, P., Robinson, O., & Roiser, J. (2026). State-, Trait- and Recovery-Related Differences in Reinforcement Learning and Value-Based Decision Making in Major Depressive Disorder. eLife, 15. 10.7554/eLife.112619.1

O’Reilly, J. X., Woolrich, M. W., Behrens, T. E. J., Smith, S. M., & Johansen-Berg, H. (2012). Tools of the trade: Psychophysiological interactions and functional connectivity. Social Cognitive and Affective Neuroscience, 7(5), 604~609. 10.1093/scan/nss055

Poldrack, R. A. (2007). Region of interest analysis for fMRI. Social Cognitive and Affective Neuroscience, 2(1), 67~70. 10.1093/scan/nsm006

Sazhin, D., Dachs, A., & Smith, D. V. (2025). Meta-Analysis Reveals That Explore–Exploit Decisions Are Dissociable by Activation in the Dorsal Lateral Prefrontal Cortex, Anterior Insula, and Dorsal Anterior Cingulate Cortex. European Journal of Neuroscience, 61(6), e70081. 10.1111/ejn.70081

Shenhav, A., Botvinick, M. M., & Cohen, J. D. (2013). The Expected Value of Control: An Integrative Theory of Anterior Cingulate Cortex Function. Neuron, 79(2), 217~240. 10.1016/j.neuron.2013.07.007

Timashkov, A., Anderson, S., & Zinchenko, O. (2025). Neural correlates of uncertainty processing: Meta-analysis of fMRI studies. Frontiers in Neuroscience, 19. 10.3389/fnins.2025.1662272

Todd, J. J., & Marois, R. (2004). Capacity limit of visual short-term memory in human posterior parietal cortex. Nature, 428(6984), 751~754. 10.1038/nature02466

Turker, S., Gerardos, G., Büch, F., Fumagalli, B., Kuhnke, P., & Hartwigsen, G. (2026). Task load modulates network interactions between bilateral fronto-parietal and cerebellar areas during verbal working memory. iScience, 29(5), 115502. 10.1016/j.isci.2026.115502

Valdebenito-Oyarzo, G., Martínez-Molina, M. P., Soto-Icaza, P., Zamorano, F., Figueroa-Vargas, A., Larraín-Valenzuela, J., Stecher, X., Salinas, C., Bastin, J., Valero-Cabré, A., Polania, R., & Billeke, P. (2024). The parietal cortex has a causal role in ambiguity computations in humans. PLOS Biology, 22(1), e3002452. 10.1371/journal.pbio.3002452

Wan, Q., & Sloutsky, V. (2026). Working Memory Shapes Information Sampling and Attention Allocation Across Development. Journal of experimental psychology. General, 155(2), 479~498. 10.1037/xge0001848

Wiehler, A., Chakroun, K., & Peters, J. (2021). Attenuated Directed Exploration during Reinforcement Learning in Gambling Disorder. The Journal of Neuroscience, 41(11), 2512~2522. 10.1523/JNEUROSCI.1607-20.2021

Wilson, R. C., Geana, A., White, J. M., Ludvig, E. A., & Cohen, J. D. (2014). Humans Use Directed and Random Exploration to Solve the Explore–Exploit Dilemma. Journal of experimental psychology. General, 143(6), 2074~2081. 10.1037/a0038199

Witte, K., Thalmann, M., & Schulz, E. (2025). Model-based exploration is measurable across tasks but not linked to personality and psychiatric assessments. Scientific Reports, 15(1), 27479. 10.1038/s41598-025-09152-2

Worsley, K. J., Marrett, S., Neelin, P., Vandal, A. C., Friston, K. J., & Evans, A. C. (1996). A unified statistical approach for determining significant signals in images of cerebral activation. Human Brain Mapping, 4(1), 58~73. 10.1002/(SICI)1097-0193(1996)4:1%3C58::AID-HBM4%3E3.0.CO;2-O

Wyatt, L. E., Hewan, P. A., Hogeveen, J., Spreng, R. N., & Turner, G. R. (2024). Exploration versus exploitation decisions in the human brain: A systematic review of functional neuroimaging and neuropsychological studies. Neuropsychologia, 192, 108740. 10.1016/j.neuropsychologia.2023.108740

Zhao, W., Su, K., Zhu, H., Kaiser, M., Fan, M., Zou, Y., Li, T., & Yin, D. (2024). Activity flow under the manipulation of cognitive load and training. NeuroImage, 297, 120761. 10.1016/j.neuroimage.2024.120761

