## Supplementary Information for "Cognitive Load modulates the exploration-exploitation trade-off Through Reconfiguration of an IPS-Centered Decision Network"

**Supplementary method 1**

To quantify how cognitive load affected the exploration–exploitation trade-off in a dynamic reward environment, we fitted participants' trial-by-trial choices using five reinforcement-learning models (Daw et al., 2006; Wiehler et al., 2021). The models differed in how they learned option values and how expected value, uncertainty, and choice perseveration influenced action selection. In Experiment 1, the five models were fitted and compared separately for the high- and low-load conditions. The best-fitting model was then applied to both load conditions in Experiment 2 without repeating model selection.

We first implemented a Q-learning model. Let $c_t$ denote the selected option on trial $t$, $r_t$ the corresponding reward, and $Q_{i,t}$ the expected value of option $i$ before the current reward was observed. The reward prediction error for the chosen option was

$$\delta_t = r_t - Q_{c_t,t},$$

and the expected value of that option was updated according to

$$Q_{c_t,t+1} = Q_{c_t,t} + \alpha \delta_t,$$

where $\alpha$ is a constant learning rate ($0 \le \alpha \le 1$). The values of unchosen options were unchanged. Choice probabilities were calculated with a softmax function:

$$P(c_t = i) = \frac{\exp(\beta Q_{i,t})}{\sum_{j=1}^{4} \exp(\beta Q_{j,t})}.$$

Here, $\beta$ is the inverse temperature, which controls the sensitivity of choices to differences in estimated option value. Higher $\beta$ indicates more consistent selection of high-value options, whereas lower $\beta$ indicates greater choice stochasticity. Because Q-learning tracks expected values but not their uncertainty, this model does not explicitly represent uncertainty-directed exploration.

We next implemented four Bayesian learning models based on a Kalman filter. In the dynamic four-armed bandit task, the expected reward of each option changes over time according to a Gaussian random walk. The Kalman filter tracks both the estimated mean reward $\hat{\mu}_{i,t}$, and its uncertainty $\hat{\sigma}_{i,t}$, for each option. Before a new choice, the predicted mean and variance of each option were updated to account for changes in the reward environment:

$$\hat{\mu}_{i,t}^{pre} = \lambda \hat{\mu}_{i,t-1}^{post} + (1-\lambda)\theta,$$

$$(\hat{\sigma}_{i,t}^{pre})^2 = \lambda^2 (\hat{\sigma}_{i,t-1}^{post})^2 + \sigma_d^2.$$

The decay parameter $\lambda$ determines the persistence of the previous value estimate, $\theta$ is the mean-reversion center, and $\sigma_d$ is the standard deviation of the diffusion noise, these parameters were fixed at $\lambda = 0.9836$, $\theta = 50$, and $\sigma_d = 2.8$. The observation-noise standard deviation was also fixed at $\sigma_o = 4$. These quantities were treated as fixed properties of the learning model, not as load-dependent free choice parameters.

After choosing option $c_t$ and receiving $r_t$, the model calculated a prediction error and an uncertainty-dependent learning rate (the Kalman gain):

$$\delta_t = r_t - \hat{\mu}_{c_t,t}^{pre},$$

$$\kappa_t = \frac{\left(\hat{\sigma}^{pre}_{c_t,t}\right)^2}{\left(\hat{\sigma}^{pre}_{c_t,t}\right)^2 + \sigma_o^2}.$$

The posterior mean and variance for the chosen option were updated as

$$\hat{\mu}^{post}_{c_t,t} = \hat{\mu}^{pre}_{c_t,t} + \kappa_t \delta_t,$$

$$\left(\hat{\sigma}^{post}_{c_t,t}\right)^2 = \left(1 - \kappa_t\right)\left(\hat{\sigma}^{pre}_{c_t,t}\right)^2.$$

For options that were not chosen, no reward-dependent correction was applied on that trial. Accordingly, sampling an option updates its expected value and reduces uncertainty, whereas uncertainty about unchosen options evolves with the random-walk process. Unlike the constant learning rate in Q-learning, the Kalman gain changes from trial to trial as a function of the current uncertainty and observation noise.

The four Kalman-filter models used the same learning algorithm but different choice rules. In the simplest version, choices were determined only by estimated mean reward through a softmax rule, $P\left(c_t = i\right) \propto \exp\left(\beta \hat{\mu}^{pre}_{i,t}\right)$. To capture directed exploration, a second model added an uncertainty bonus $\varphi \hat{\sigma}^{pre}_{i,t}$ to each option's decision value. The parameter $\varphi$ determines the influence of uncertainty on choice: larger values indicate a greater tendency to favor uncertain options. A third model included a perseveration bonus $\rho 1\left(c_{t-1} = i\right)$, where the indicator equals 1 when option $i$ was chosen on the preceding trial and 0 otherwise. Positive $\rho$ favors repeating the preceding choice, whereas negative $\rho$ favors switching. The fourth, full Kalman-filter model included both bonuses. Its decision value was

$$V_{i,t} = \hat{\mu}^{pre}_{i,t} + \varphi \hat{\sigma}^{pre}_{i,t} + \rho 1\left(c_{t-1} = i\right),$$

with choice probability

$$P\left(c_t = i\right) = \frac{\exp\left(\beta V_{i,t}\right)}{\sum_{j=1}^{4} \exp\left(\beta V_{j,t}\right)}.$$

Setting $\varphi = 0$, $\rho = 0$, or both to zero yielded the corresponding reduced Kalman-filter models. The complete model space therefore comprised (1) Q-learning with softmax, (2) Kalman filter with softmax, (3) Kalman filter with softmax and uncertainty bonus, (4) Kalman filter with softmax and perseveration bonus, and (5) Kalman filter with softmax, uncertainty bonus, and perseveration bonus. The designation three-parameter Kalman model refers to the three choice-related parameters $\beta$, $\varphi$, and $\rho$; it does not count the fixed random-walk parameters.

Finally, the trialwise estimates from the fitted model in Experiment 2 were used to classify choices for the subsequent fMRI analysis, following the operational approach of Wiehler et al. (2021). A trial was classified as exploitation if the chosen option had the highest combined decision value, including expected value, uncertainty bonus, and perseveration bonus (equivalently, the highest softmax choice probability). Trials on which another option was selected were classified as exploration. Among exploration trials, choices of the option with the largest model-based uncertainty bonus were classified as directed exploration (DE), and the remaining trials as random exploration (RE).

**Supplementary result 1**

After excluding the 10 participants who did not perform any DE trials, the same pattern observed in Experiment 1 and in the full sample of Experiment 2 before exclusion was still observed: phi was significantly higher under low cognitive load than under high cognitive load ($t = 5.3808$, $p < 0.0001$, supplementary figure 1b). We also found that beta was significantly higher under low cognitive load ($t = 4.1906$, $p = 0.0001$, supplementary figure 1a).

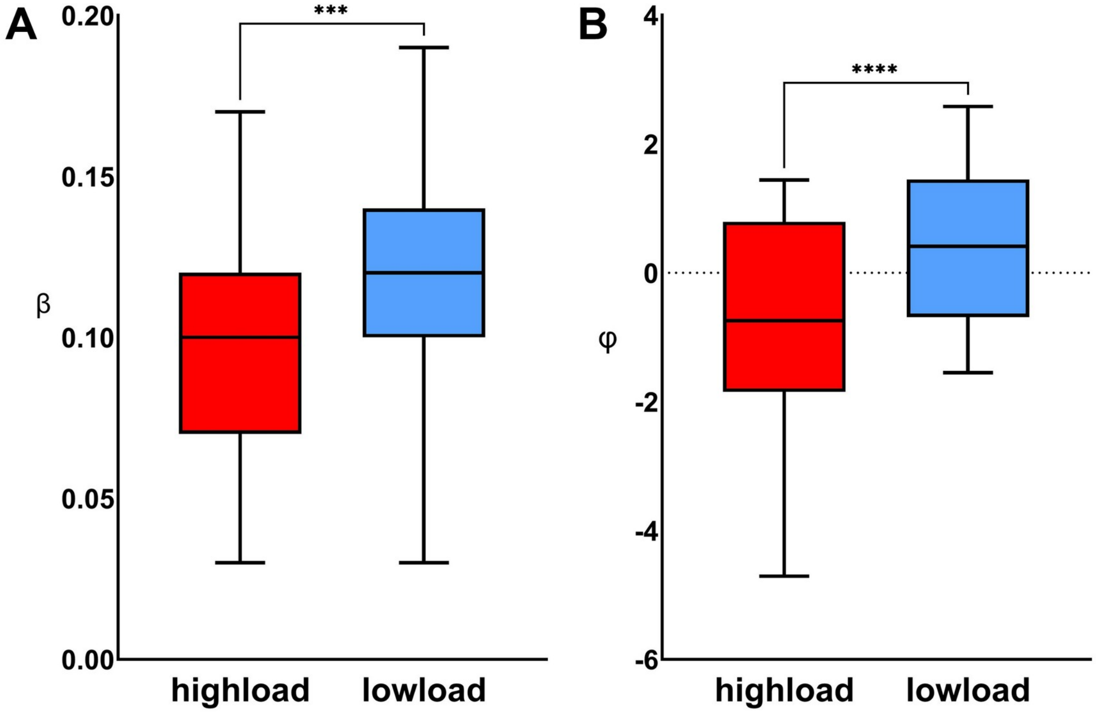


Figure 1. Behavioral model results in Experiment 2 fMRI analysis. (A) Comparison of the exploitation parameter beta between high- and low-load conditions. beta was significantly higher under low cognitive load. (B) Comparison of the exploration bonus parameter (φ) between high- and low-load conditions. φ was significantly higher under low cognitive load. ***, $p < 0.001$, ****, $p < 0.0001$.

***References***


Daw, N. D., O’Doherty, J. P., Dayan, P., Seymour, B., & Dolan, R. J. (2006). Cortical substrates for exploratory decisions in humans. *Nature*, *441*(7095), 876〜879. https://doi.org/10.1038/nature04766

Wiehler, A., Chakroun, K., & Peters, J. (2021). Attenuated Directed Exploration during Reinforcement Learning in Gambling Disorder. *The Journal of Neuroscience*, *41*(11), 2512〜2522. https://doi.org/10.1523/JNEUROSCI.1607-20.2021
